# Deep Learning Unravels Potential Antibiotic-Resistance Drugs against Klebsiella Pneumoniae

**DOI:** 10.1101/2024.11.25.625191

**Authors:** Peng Lin, Jiali Zhou, Sijun Meng, Zhihao Lei, Linxiao Jin, Hesong Qiu, Yingfan Xu, Tsehao Hsu, Yemei Bu, Glen Qin, Wen Zhang

## Abstract

**Background:** Klebsiella pneumoniae (KP) is a critical pathogen recognized by the World Health Organization due to its high mortality rates and resistance to antibiotics. Developing effective treatments for KP is essential to mitigate the growing threat of multidrug-resistant infections. This study leverages deep learning techniques to identify and validate potential drug candidates against KP, accelerating the drug discovery process and reducing associated risks.

**Results:** A comprehensive dataset was constructed, comprising 3,475 drugs selected from the DrugBank database, filtered for their potential efficacy and safety. Using convolutional neural networks (CNNs) for drug-drug interaction analysis, a prediction accuracy of 72% was achieved. Evolutionary Scale Modeling (ESM) was employed to calculate molecular similarities between drugs and KP strains, identifying five promising candidates with structural similarities exceeding 85% to known KP targets. A drug repurposing knowledge graph (DRKG) further validated these findings, with the PairRE model outperforming other approaches in ranking potential drug candidates.

**Conclusions:** This study presents a scalable and efficient framework for identifying and validating drug candidates to combat antibiotic-resistant KP. By integrating deep learning models, molecular similarity analysis, and knowledge graph techniques, this approach shortens the drug discovery timeline from years to months. The identified candidates hold significant promise for preclinical and clinical testing, offering a pathway to effective therapies against KP and other multidrug-resistant pathogens.

## 1 Introduction

Klebsiella Pneumonia (KP), a WHO [1] priority pathogen, is a bacterial strain that responsible for diverse infections, often resulting in high fatality rates. In cases of antibiotic-resistant (drug resistance) KP, patients endure prolonged severe chest pain, difficulty of breathing, fever, and cough with reddish jelly-like mucus, posing a significant risk of death as reported by [2, 3]. The epidemiology and resistance patterns of common intensive care unit infections was evaluated by [3], noting Klebsiella as the most common causative organism, with multi-drug resistance observed in 75.5% of cases. The clinical characteristics, risk factors, and outcomes of KP pneumonia were investigated by [4] along with bloodstream infections. Hence, effective drug development is an imperative health priority to treat KP infections.

To propel the discovery of antimicrobial drugs against KP, deep learning (DL) methods were employed and described herein. Clinical gene sequence data were gathered and studied by [5] from over 3,000 patients diagnosed with KP bacteria, providing insights into the genetic aspects of the infection’s impact on health. Concluding remarks and recommended actions are drawn to drive further research and development in combating KP infections.

Through genetic experiments and genomic analysis, [6] investigated the transfer, stability, and cargo genes linked with mobilizable plasmids in KP, aiming to elucidate their role in horizontal gene transfer. Their study concluded that mobilizable plasmids play a significant role in facilitating the dissemination of antimicrobial resistance and virulence genes in KP, potentially enabling co-transfer alongside conjugative plasmids while evading CRISPR-Cas system restrictions.

Mobile colistin resistance (mcr) variants in KP isolates from humans were identified by [5] in China, detecting six variants associated with diverse plasmid groups. They also analyzed Multiplexed Intermixed CRISPR (MICs) for mcr-positive isolates, revealing mcr-10.1 susceptibility, and highlighted mcr transmission between humans and the environment. The transfer, stability, and cargo genes of the mobilizable plasmids of KP were examined via genetic experiments and genomic analysis. Deep Learning (DL) has proven to be a valuable asset in drug discovery, offering an insightful perspective on the current state-of-the-art in this field. This approach not only sheds light on existing challenges but also highlights successful applications and opportunities for further advancements. The synergy of advanced hardware and algorithms has contributed to the widespread adoption of machine learning, particularly DL, in drug discovery. Recent comprehensive overviews by [7-10] provide a detailed exploration of the evolving landscape.

DL has been successfully applied in drug discovery as detailed view of the current state-of-the-art in drug discovery and highlighting not only the problematic issues, but also the successes and opportunities for further advances. Amid the COVID-19 crisis, [11] compiled a vast database of experimentally validated drug-virus associations through text mining, developed a novel WRMF approach integrating known drug-virus associations, drug-drug chemical structure similarity, and virus-virus genomic sequencing similarity networks, outperforming state-of-the-art methods. WRMF demonstrates accuracy and reliability in identifying potential drugs for novel viruses, providing a valuable tool for drug repurposing efforts. In [9] study, a novel data-driven approach was introduced to identify assay interferents and prioritized true bioactive compounds from high throughput screening data. The method distinguishes between compounds with desired biological responses and assay artifacts without prior screens or interference mechanisms. It consistently outperforms tested baselines, requires less than 30 seconds per assay on low-resource hardware, and is ideal for guiding pharmacological optimization post high throughput screening.

A CNN architecture was constructed by [12] with 5 layers and 3 × 1 kernel size to predict drug–drug interactions to first extract feature interactions from drug categories, targets, pathways and enzymes as feature vectors and then use the Jaccard similarity as the measurement of drugs similarity. The concluded result showed CNN with multiple drug features is more effective than single feature on predicting drug-drug interactions (DDIs).

As pointed out by [13], AI-based computing programs can significantly accelerate the drug design and development process, completing discovery to preclinical stages in less time than the industry average of five to six years. Venture-funded companies have reported timelines of fewer than 18 months from target identification to candidate identification.

Relevant DL applications and case studies were analyzed by [7], providing a comprehensive view of drug discovery’s current state and highlighting challenges, successes, and opportunities. AI technology holds potential in addressing key obstacles by leveraging in silico approaches for de novo drug design, synthesis prediction, and bioactivity prediction, thereby reducing costs, time, and workloads in early-stage drug discovery.

DL methods of deep neural networks (DNN), convolutional neural networks (CNN), and multi-task learning (MTL) and their effectiveness were studied by [8] in repurposing existing drugs for COVID-19 treatment, like remdesivir and oseltamivir, highlighting its potential in drug discovery. A multimodal deep learning framework named DDIMDL that combines diverse drug features was proposed by [14] to build a model for predicting DDI-associated events. DDIMDL achieves an accuracy of about 88% and an area under the precision–recall curve of about 92%.

A study by [15] demonstrates that combining SERS spectroscopy with a CNN-attention mechanism achieved a high prediction accuracy of 99.46% in identifying carbapenem-sensitive, carbapenem-resistant, and polymyxin-resistant Klebsiella pneumoniae strains. This approach highlights the potential for rapid and accurate clinical antibacterial susceptibility testing. Additionally, mobilizable MDR and virulence plasmids of Klebsiella pneumoniae were studied by [6]. Their findings indicate that these plasmids can transfer with helper conjugative CR plasmids, evade CRISPR-Cas defenses, and promote the spread of antibiotic resistance and virulence genes. These results underscore the significant threat posed by mobilizable plasmids in the dissemination of these harmful traits.

Aside from literature review of related work in the field, current paper presents a detailed investigation into how drug candidates combating KP virus are identified and validated, organized into sections covering DL methods, data extraction, molecular similarity comparison using EMS Model and WRMF, and the DRKG approach.

## 2 Materials and Methods

### 2.1 Datasets

Three primary datasets, as shown in Figure 1, are utilized for this investigation. They are

**Figure 1.**
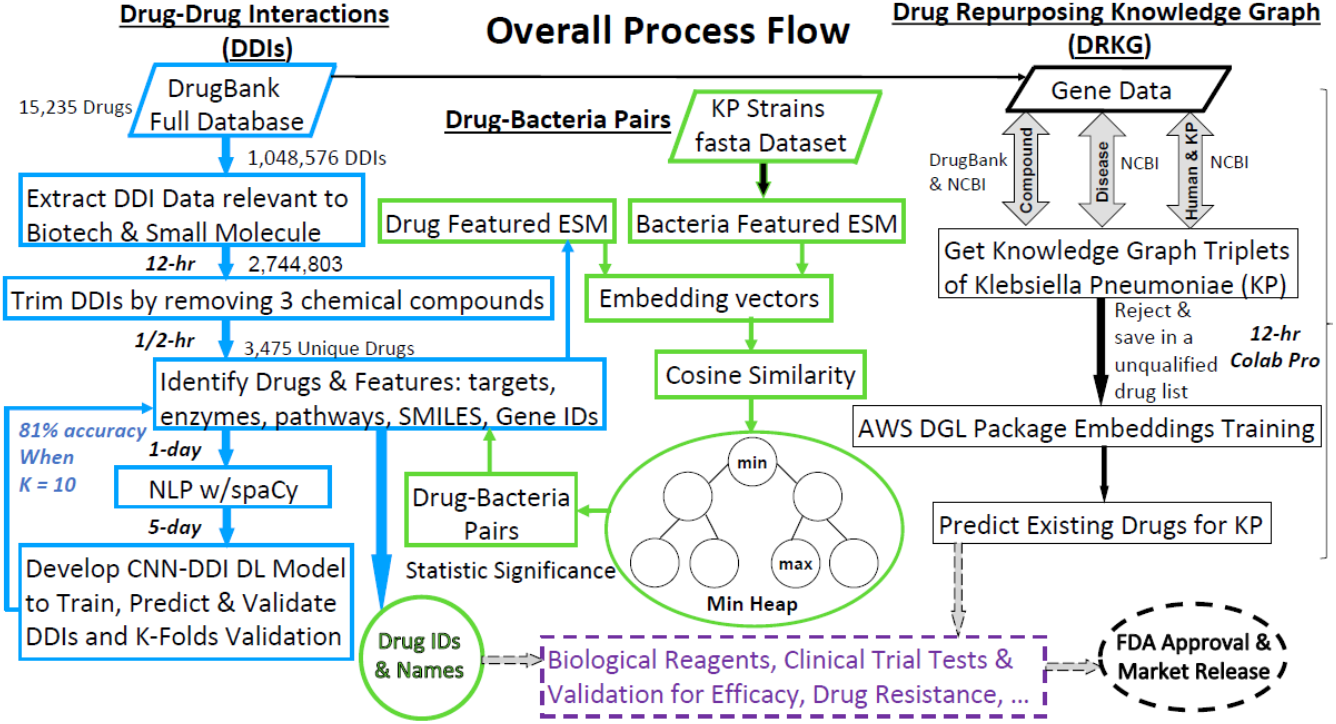
Overall Process of DDIs and Unique Drugs Extraction, CNN, ESM with Drug-Bacteria Similarity, and DRKG. ***Left***: *DrugBank database for extracting low-risk DDIs, selecting unique drugs with target proteins and training, predicting, and validating using NLP and CCN/DDI deep learning*. ***Middle***: *KP virus strain FASTA dataset for ESM-based similarity analysis between drug targets and KP strains, identifying top drug candidates using a mini heap alogrithm*. ***Right***: *NCBI database (GeneBank and PubMed) for drug repurposing knowledge graph analysis*.

1. DrugBank full database: a comprehensive database [16] containing a unique bioinformatics and cheminformatics resource on drugs and drug targets created and maintained by the University of Alberta in Canada. This full database is used for extracting drug-drug interactions with low risk and no adverse effects for identifying potential unique drugs with key features of Targets, Enzymes, SMILES, Pathways. Year 2023 version of full database file was used.
2. KP strain fasta dataset [5]: consists of 831 virus strains collected from patients in hospitals in China. The fasta dataset is used to extract target proteins for similarity analysis between potential drugs and KP virus strains.
3. National Center for Biotechnology Information (NCBI) databases: include GenBank for DNA sequences and PubMed used for Drug repurpose knowledge graph analysis and characterization.

### 2.2 Data Extraction

DrugBank full database was first used to filter out unwanted drug types: Vaccine, Antibody, Toxin, Biological, Biopolymer, Oligonucleotide, and Oligosaccharide and to secure two types of biotech and small molecule drugs as shown in Figure 2. 2,744,803 drug pairs of DDIs were then extracted covering risk and effects of drug-drug interactions. A partial list of effects of drug-drug interactions, along with their actions, is shown in Table 1 when two drugs are combined or used together.

**Table 1.**
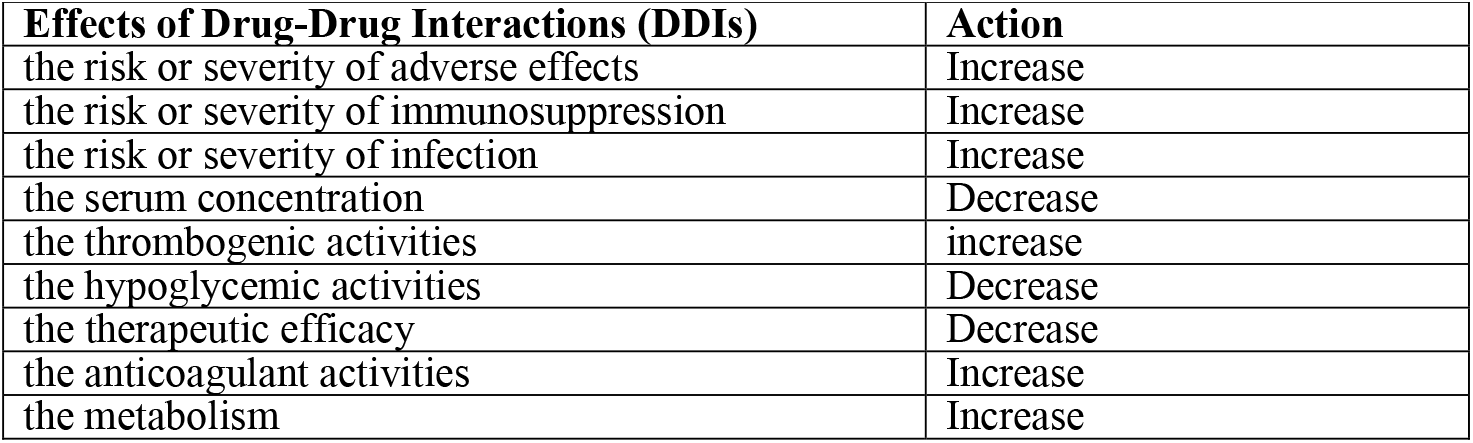
Effects of drug-drug interactions when 2 drugs are combined.

**Figure 2.**
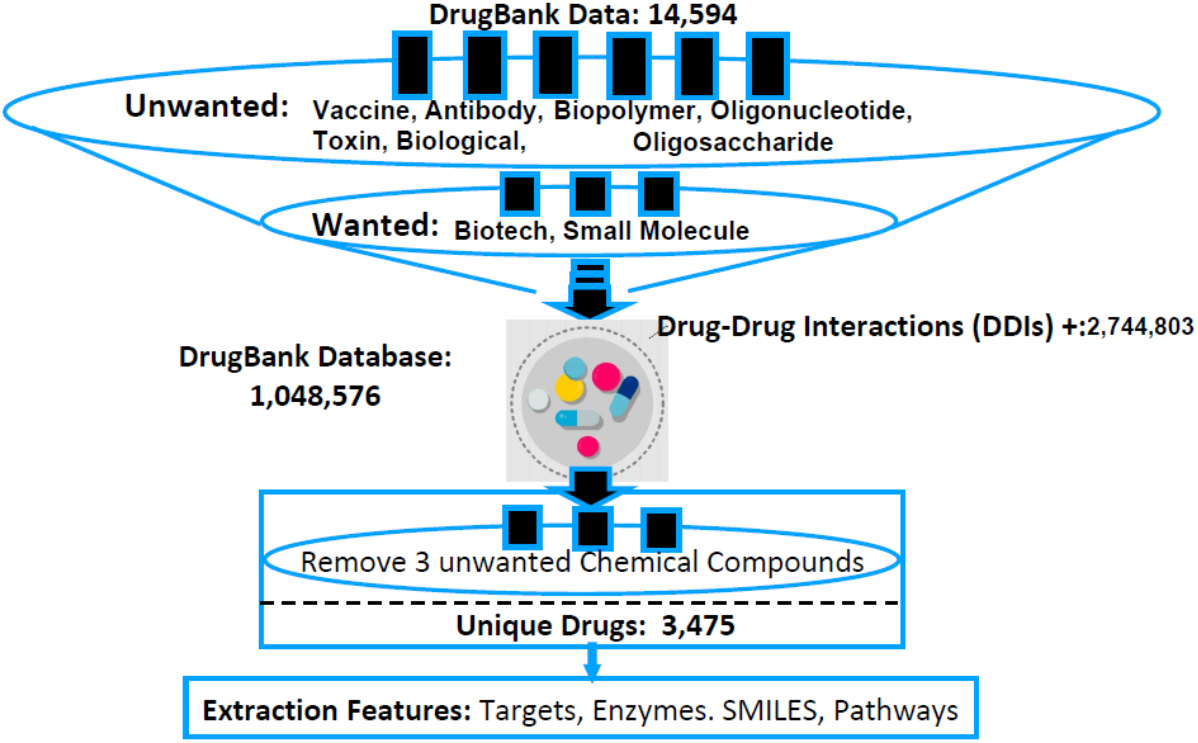
Process Flow of DDI & Unique Drugs by removing unwanted types and chemical compounds. ***Unique drugs w/related targets from biotech and small molecule types***

Three chemical compounds, irrelevant to potential drugs to combat KP bacteria, get removed. Those compounds are

1. (4-{(2S)-2-[(tert-butoxycarbonyl)amino]-3-methoxy-3-oxopropyl}phenyl)methaneseleninic acid: methaneseleninic acid is an organoselenium compound, a seleninic acid with the chemical formula CH3SeO2H.
2. (S)-2-Amino-3-(4h-Selenolo[3,2-B]-Pyrrol-6-Yl)-Propionic Acid: Propionic acid is a naturally occurring carboxylic liquid acid with a pungent and unpleasant smell resembling body odor with chemical formula C3H6O2
3. Florbetaben (18F): is a diagnostic radio-tracer as an imaging agent developed for routine clinical application to visualize beta amyloid plaques in the brain via a PET scan. Its chemical formula is C21H2618FNO3.

At the end of extraction processes, 3,475 unique drugs after removing any duplication of drug pairs and three unwanted chemical compounds and/or elements.

Critical features of targets, enzymes, SMILEs, and pathways are critical features for drug discovery. Basic definitions of these features are reviewed as follows:

Targets: refer to specific biological molecules, like proteins or nucleic acids associated with diseases. Drugs interact with these targets to achieve therapeutic effects.

Enzymes: are proteins that catalyze biochemical reactions in living organisms. Drugs targeting enzymes regulate or block their activity, altering disease-related processes r profession.

SMILES: is a textual representation of a molecule’s chemical structure. Used for computer-based modeling to predict properties and interactions.

Pathways: refer to sequences of biochemical reactions in cells or organisms to perform specific biological functions. They represent interconnected networks of molecules and processes that are dysregulated in disease states. Drugs can be designed to target specific pathways to restore normal physiological function or disrupt disease-related processes.

To extract data features about Targets, Enzymes, Pathways, and SMILES for relevant drugs from the DrugBank full dataset, following steps were carried out.

Upon gaining an approval from DrugBank, the full dataset file in XML format was downloaded for this work as described in this section by one of the authors who had signed an NDA.

By parsing the DrugBank dataset, the relevant features were extracted containing information about drugs with IDs and names, including their targets, enzymes, pathways, and SMILES representations. Pathways associated with each drug in the dataset were located. Extract pathway names, small molecular identifiers, and any relevant information about their biological significance or relevance to drug action. SMILES (Simplified Molecular Input Line Entry System) is a compact notation used to represent the structure of chemical compounds critical to the analysis. Thus, the SMILES representation was extracted for each drug molecule.

These five key features are extracted for subsequent CNN data training and validation and ESM final drugs screening and selections.

Overall, understanding the role of targets, enzymes, SMILES, and pathways in drug discovery is crucial for designing and developing effective therapeutic interventions to treat various diseases and conditions. These features provide valuable insights into the molecular mechanisms underlying diseases and help guide the rational design of new drugs with improved efficacy and safety profiles.

### 2.3 NLP and CNN/DDI

Natural language processing (NLP) is used to process a dataset of drug-drug interactions (DDIs) and to extract key interaction information. For each interaction sentence, spaCy’s NLP pipeline is used to analyze the sentence structure, identify the root action, determine the mechanism of the drug interaction, action verb (increase or decrease), and the two drugs involved in the interaction.

Convolutional Neural Networks (CNNs) and Drug-Drug Interactions (DDIs) analysis were combined for deep learning training and validation to assess drug candidates with reasonable accuracy. As aforementioned dataset containing features of Targets, Enzymes, Pathways, and SMILES about 3,475 drugs, preprocessing and encoding the data into a format suitable for input into the CNN model.

The CNN model comprises 9 layers: 2 convolutional, 2 batch normalization, 1 max pooling, 1 flatten, 2 dense, and 1 output layer, as shown in Figure 3. The convolutional layers extract spatial features from input data, which are processed by dense layers. ReLU activation in convolutional and dense layers introduces non-linearity, while the softmax function in the output layer handles multi-class classification. The model is trained using the Adam optimizer with a learning rate of 0.001 and categorical cross-entropy loss to classify 50 drug-drug interaction effects.

**Figure 3.**
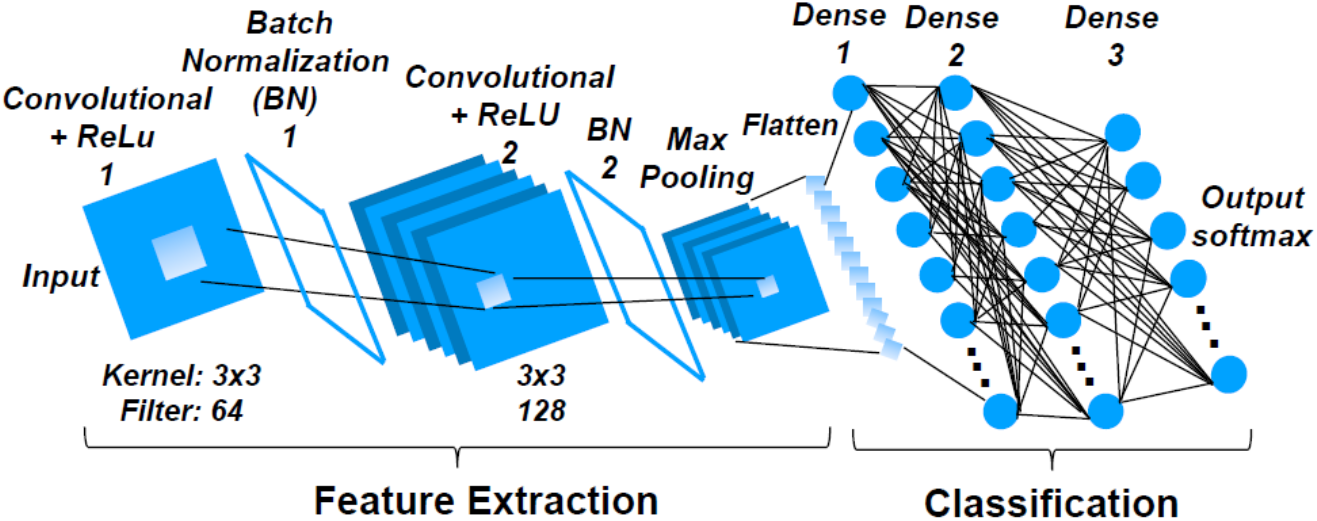
Convolutional Neural Networks with 9 Layers. ***1st layer:*** *Input (Convolutional)*, ***2nd*** *layer: Batch Normalization*, ***3rd***: *Convolutional*, ***4th***: *Batch Normalization*, ***5th***: *Max Pooling*, ***6th***: *Flatten*, ***7th***: *Dense*, ***8th***: *Dense*, ***9th***: *Output (Dense)*

Hyperparameters such as the learning rate, number of filters in convolutional layers, batch size, and number of epochs were fine-tuned to achieve optimal performance. Regularization techniques, including dropout and batch normalization, prevent overfitting and enhance generalization. This approach helps understand drug interactions’ impacts on the immune system, metabolism, and circulation, supporting effective treatment and reducing side effects.

The CNN model is trained on prepared datasets to identify patterns and relationships between drugs using features like SMILES, Targets, and DDI labels. During training, hyperparameters are adjusted iteratively to minimize the difference between predicted and actual DDI labels. Model performance is then evaluated on a validation dataset, measuring metrics such as accuracy, precision, and recall to assess its ability to generalize to unseen data.

If needed, the model undergoes fine-tuning through hyperparameter adjustments, architecture modifications, or data augmentation to enhance performance. Once validated, the model is deployed for practical use, with ongoing monitoring to ensure effectiveness and reliability.

By leveraging CNNs for DDI analysis, machine learning significantly accelerates drug discovery, repurposing, and pharmacovigilance com-pared to traditional methods.

### 2.4 Molecular Similarity Comparison

In this study, we utilized Evolutionary Scale Modeling (ESM) [17], a set of transformer protein language models developed by the Meta Fundamental AI Research Protein Team (FAIR). The ESM-2 model, specifically, has shown superior performance compared to all tested single-sequence protein language models across a range of structure prediction tasks. ESMFold, a model harnessing ESM-2, generates accurate structure predictions directly from the sequence of a protein.

We leveraged ESMFold to extract normalized embedding vectors for both the drugs in DrugBank[citation needed] and the target proteins of 831 strains of KP. These normalized vectors facilitated a direct and unbiased comparison through cosine similarity measures, allowing for an accurate assessment of molecular resemblance. Thus, Figure 4 depicts the process flow of ESM to derive drug-bacteria similarity from comparing two datasets: 831 bacterial strains vs. 3,475 drugs with features of targets and others.

**Figure 4.**
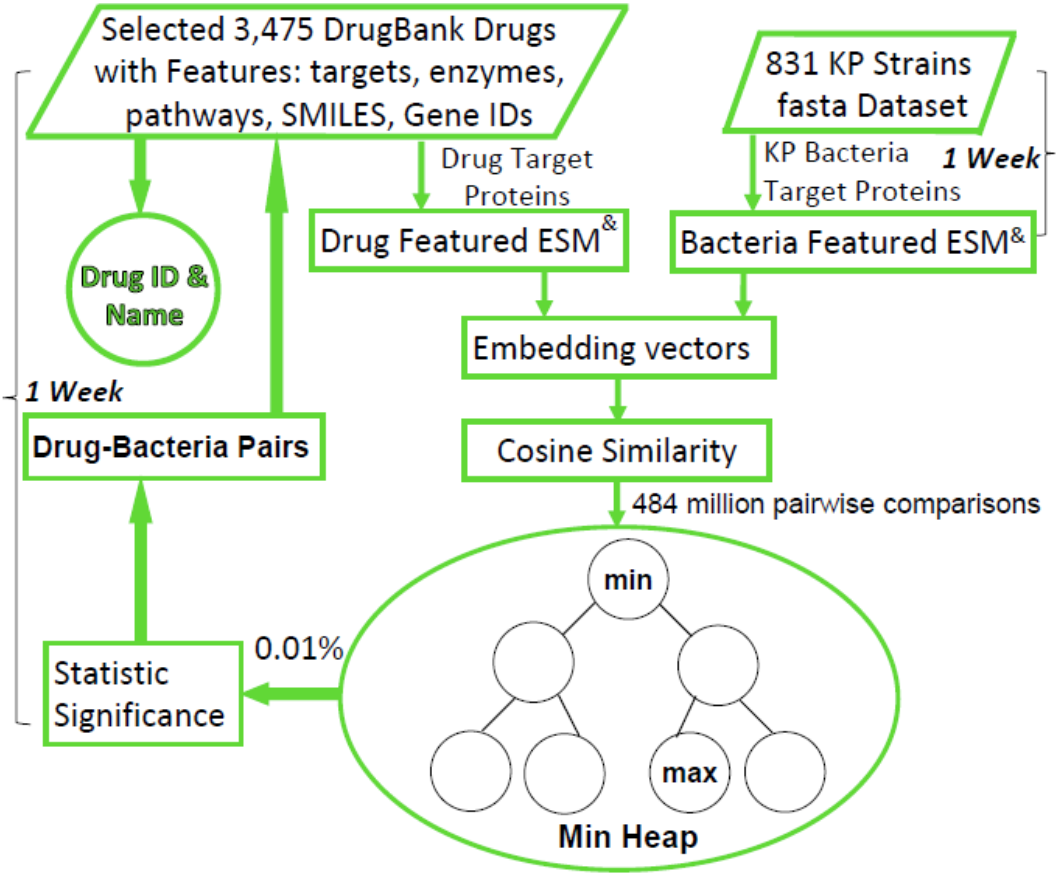
Process Flow of Drug-Bacterial Pairs Similarity Analysis. ***ESM normalized embedding vectors for cosine similarity calculations to determine structural similarities between drugs and KP bacteria using min heap algorithm for identifying top drug candidates***.

**Figure 5.**
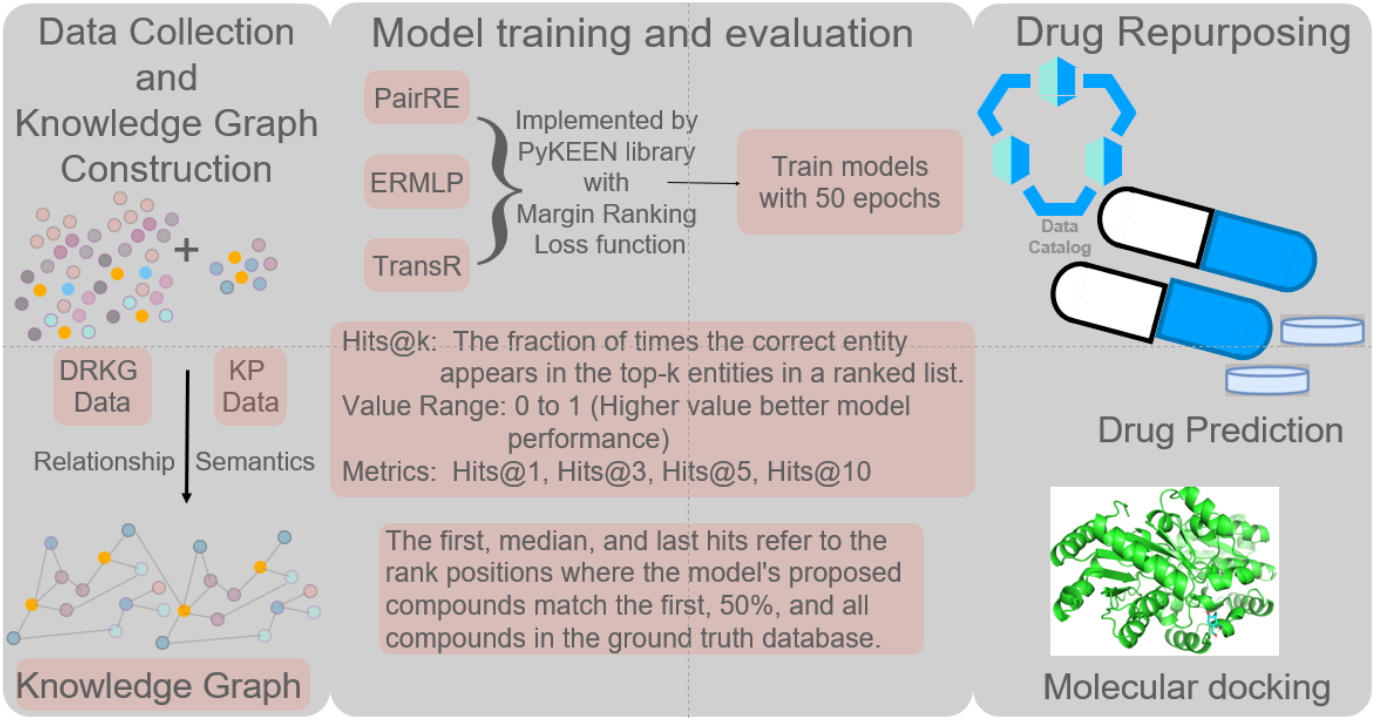
DRKG Process from Data Collection to Drug Repurposing

The ESM model’s ability to generate highly detailed embeddings for proteins and compounds provided us with a robust foundation for examining the structural similarities. The embeddings were essential for assessing the potential of drugs to interact with various bacterial strains. By focusing on KP, a pathogen associated with significant clinical challenges, this approach aimed to pinpoint potential therapeutic agents that might be effective in treating infections caused by this bacterium. The use of ESM’s embeddings ensured that our analysis was rooted in cutting-edge protein modeling technology.

Upon analyzing the structural data, each drug was compared to bacterial target proteins yielding approximately 484 million pairwise comparisons. The ESM model normalized embedding vectors to ensure consistency, enabling cosine similarity calculations to determine structural similarities using the formula:

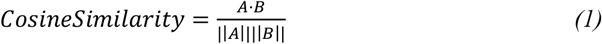

Where A and B represent two vectors being compared representing drug target proteins and bacterial molecular embeddings. ||A|| and ||B|| are their respective vector magnitudes. The cosine similarity measure, which evaluates the angular distance between two vectors, was pivotal in our analysis. It reliably gauged the resemblance between chemical structures and bacterial proteins, enabling us to quantify molecular similarity and rank potential drug candidates based on their structural alignment with bacterial targets. This approach allowed us to eliminate least promising candidates and prioritize those with higher therapeutic potential.

### 2.5 DRKG Approach

The Drug Repurposing Knowledge Graph (DRKG) complements deep learning (DL) by linking drugs, diseases, targets, and interactions through advanced graph-based algorithms It uncovers connections between drugs and diseases, facilitating the identification of new applications for exiting drugs and expediting drug repurposing. DRKG integrates data from sources like DrugBank, GNBR, Hetionet, STRING, IntAct, and DGIdb encompassing 97,238 entities across 13 types and 5,874,261 triplets across 107 edge types. Additional KP-specific data from public sources adds 211 entities and 207 triplets. A knowledge graph is constructed using association rules, and three embedding models trained on specific scoring functions predict drugs based on target relations aiding in treatments for KP and related complications.

#### 2.5.1 Validation

##### 2.5.1.1 Internal evaluation

The three knowledge graph neural network (KGNN) models were evaluated intrinsically, within the knowledge graph defined as triples and externally against clinical trial data for KP drugs to gauge real-world predictive power. Intrinsic performance was assessed using rank-based metrics, including Adjusted Mean Rank (AMR) which measures performance relative to the dataset size on a scale from 0 to 2, with lower values indicting better performance.

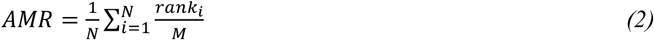

The total number of drugs in the dataset is denoted by N, the ranking of the ith real drug is represented by *rank*_*i*_ and the total number of drug candidates is designated by M.

The second indicator is the Hits at Rank k(Hits@k). Hits@k is defined as the fraction of times that the correct or “true” entity appears among the top-k entities in the ranked list. The value of hits@k is constrained between 0 and 1. The larger the value, the more accurate the model is deemed to be. For this project, I estimated the following metrics: hits@1, hits@3, hits@5, and hits@10.

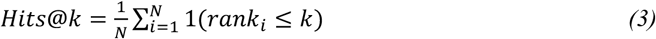

The variable N represents the total number of instances of the drug in the dataset. The variable 1 is an indicator function with a value of 1 when the condition within it holds, and 0 otherwise. The variable *rank*_*i*_ is the ranking of the correct answer in the ith instance.

##### 2.5.1.2 External evaluation

To validate the KGNN models, an external analysis compared the predicted compound rankings to drugs in clinical trials for KP, based on data from ClinicalTrials.gov. This search identified 32 trials involving 16 drugs, assessed using specific metrics:

The first hit is the ranking position at which the compounds proposed by a KGNN models match one from the ground truth database. The median hit is the rank where a KGNN model predictions match 50% of the compounds in the ground truth database. Finally, the last hit is the rank where KGNN model predictions match all the compounds from the ground truth database.

In all, smaller values for these metrics indicate better performance, as fewer predictions are needed to match real-world compounds.

## 3 Results

Results of data extraction, CNN-DDI model, ESM model with similarity analysis, and DRKG are reported herein.

### 3.1 Data Extraction

3,475 unique drugs are extracted limiting to biotech and small molecule types based on no adverse effects of DDIs. Results are illustrated in Tables 2 and 3 to show how many unique drugs were identified for each respective key feature for subsequent CNN/DDI training and validation. ESM modeling for similarity analysis to narrow down potential drug candidates based solely on the 2,155 unique drugs with the Targets feature.

**Table 2.** Drug Count with respective Features among 3,475 Drugs.

| DrugBank Drugs | Targets | Enzymes | SMILES | Pathways |
| --- | --- | --- | --- | --- |
| 3,475 | 2,155 | 889 | 3,017 | 435 |

**Table 3.**
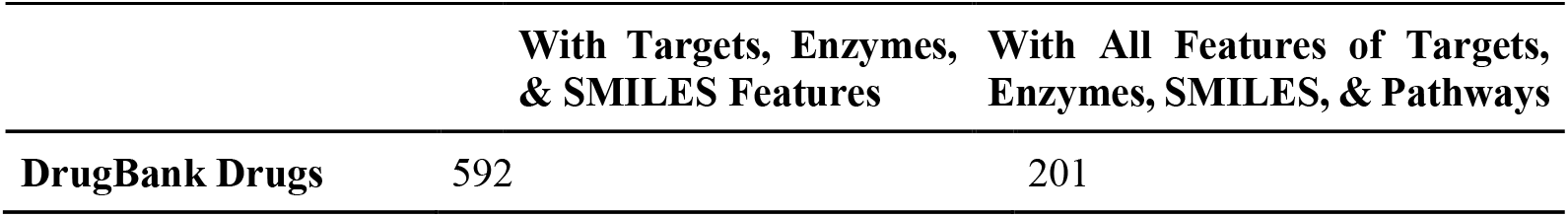
Drug Count with all Features among 3,475 Unique Drugs.

### 3.1 CNN-DDI Model

CNN/DDI model training to predict drug candidates for validation achieved an accuracy around 72% as the epoch reaches to 10 as seen in Figure 6. Moreover, K-Fold cross validation method is applied by splitting the dataset into K subsets, known as folds. The model is trained on K-1 subsets, leaving one subset for evaluation. This process is repeated K times, with a different subset reserved for testing in each iteration, leading to a consistent 75% accuracy.

**Figure 6.**
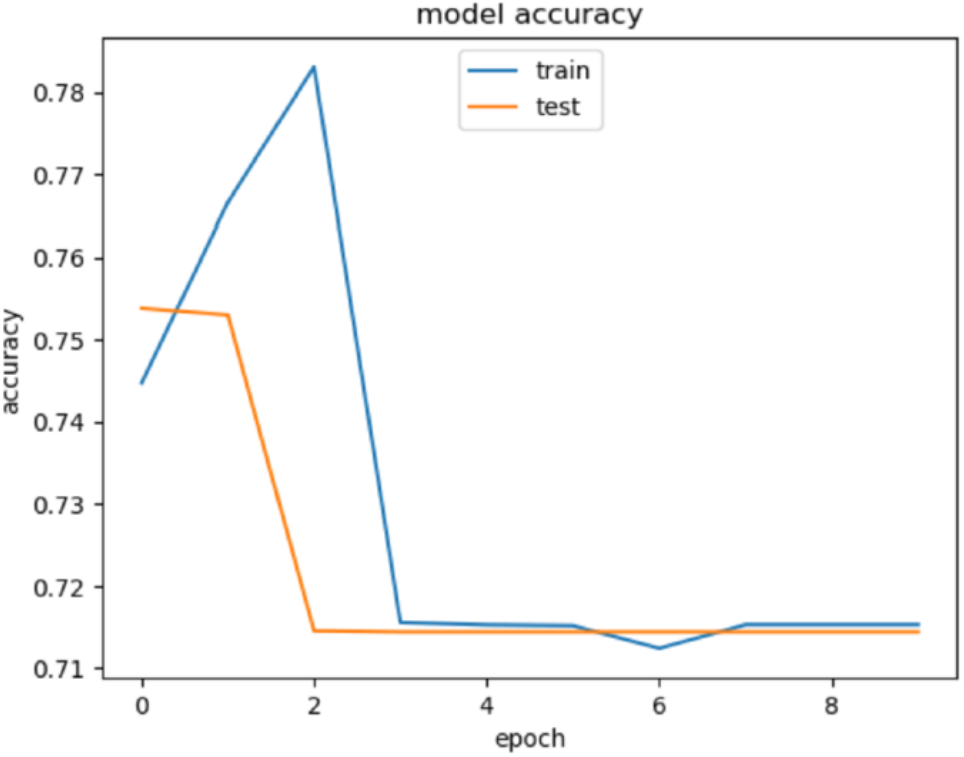
Model accuracy of Training and Validation vs. epoch

### 3.3 ESM Model with Similarity Analysis

#### 3.3.1 Molecular Similarity Comparison result

To manage the vast data from these comparisons, we utilized a priority queue, a heap-like data structure that keeps the minimum element at the top. For each strain, only the top 0.01% of pairs were retained in the queue. When encountering a new pair with higher similarity than the current top, it replaced the existing one. This method ensured the retention of the most relevant and significant comparisons, optimizing the efficiency of our analysis.

Additionally, similarities were recorded for statistical distribution analysis, revealing overall structural trends within the dataset. Figure 7 depicts these trends, showing the similaritt distribution across all drug-bacteria pairs. This approach streamlined the dataset into an accessible format while highlighting prevalent similarity ranges, offering deeper insights into common structural features.

**Figure 7.**
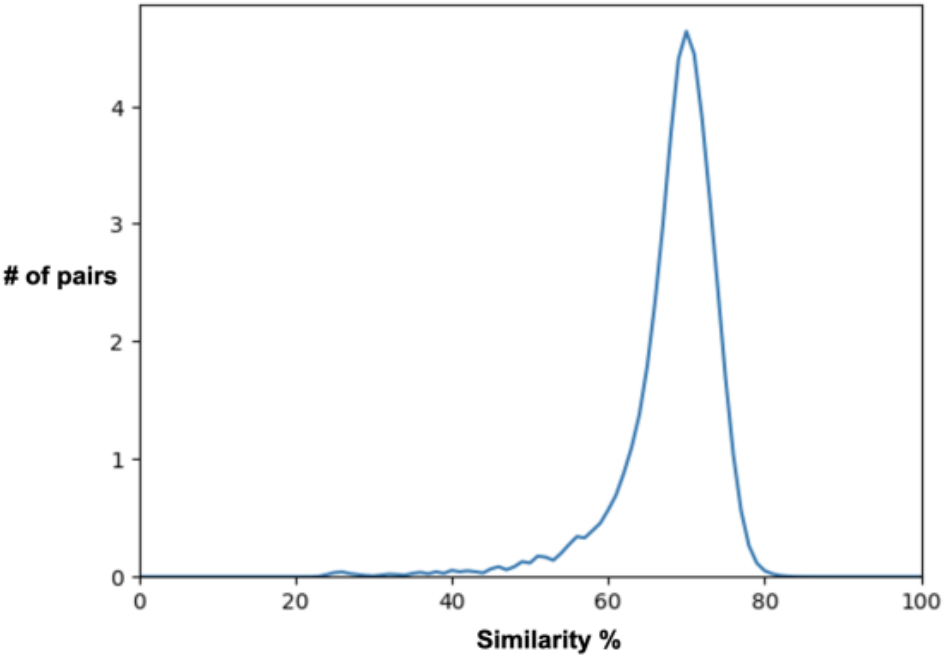
A Graph result of drug-bacterial pairs vs. % similarity

#### 3.3.2 Implications of High-Similarity Findings

Further scrutiny was applied to the top similarity scores, revealing significant matches with notable drugs such as Rofecoxib and Artenimol, which have previously been recognized for their effectiveness against KP in pharmacological studies. The identification of these drugs through our structural similarity analysis validates the effectiveness of our approach, as these drugs have been independently verified to have therapeutic potential against the bacterium. Other drugs, such as Radezolid, and elements such as copper and zinc, also featured in high-similarity pairings, albeit less frequently. These findings align well with known therapeutic effects and suggest potential avenues for exploiting these structural similarities in drug design. All drugs are small molecule type as listed in Table 4.

**Table 4.**
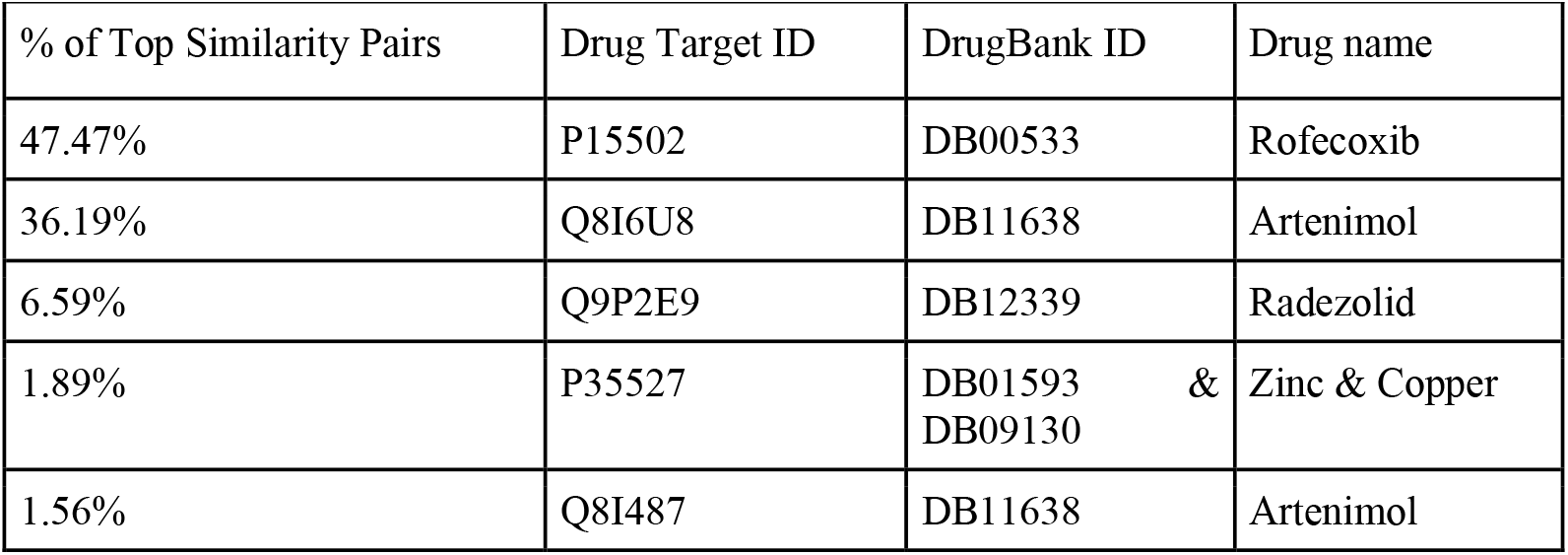
Molecular Similarity Comparison Result.

| % of Top Similarity Pairs | Drug Target ID | DrugBank ID | Drug name |
| --- | --- | --- | --- |
| 47.47% | P15502 | DB00533 | Rofecoxib |
| 36.19% | Q8I6U8 | DB11638 | Arteminol |
| 6.59% | Q9P2E9 | DB12339 | Radezolid |
| 1.89% | P35527 | DB01593 & DB09130 | Zinc & Copper |
| 1.56% | Q8I487 | DB11638 | Arteminol |

The strong correlation between high similarity scores and known therapeutic effectiveness highlights the potential of structural comparisons as a predictive tool in drug discovery. This targeted approach repurposing and development particularly valuable for combating antibiotic-resistant strains like KP. By identifying promising candidates based on molecular resemblance, this method enhances efficiency ad uncovers potential therapies that might otherwise be overlooked. High-similarity findings provide a foundation for future research and clinical trials, as structurally similar compounds are likely to exhibited strong therapeutic effects against targeted bacterial infections.

### 3.4 DRKG Results

#### 3.4.1 Internal evaluation result

As shown in Table 5, PairRE has an AMR of 0.018, the lowest of the three models, indicating that it performs the best in terms of ranking. On average, it ranks the correct drug very highly. In the case of considering only whether one of the highest-ranked drugs is correct, both PairRE and DistMult have a fairly close performance of 0.018 and 0.019, respectively. In contrast, ERMLP’s performance of 0.038 is almost twice as high as the first two. When considering whether the top three contain the correct drug, PairRE has a clear lead, with a score of 0.111, followed by ERMLP at 0.085 and DistMult at 0.049. This indicates that PairRE is more accurate in its top three predictions. In the top five, PairRE still leads (0.152), with ERMLP and DistMult at 0.115 and 0.068, respectively. When considering the top ten, the success rates of all the models increase, with PairRE at 0.218, ERMLP at 0.166, and DistMult at 0.103. This indicates that in the wider range, the PairRE still holds the lead.

**Table 5.**
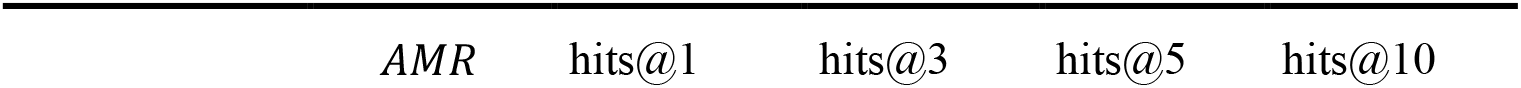
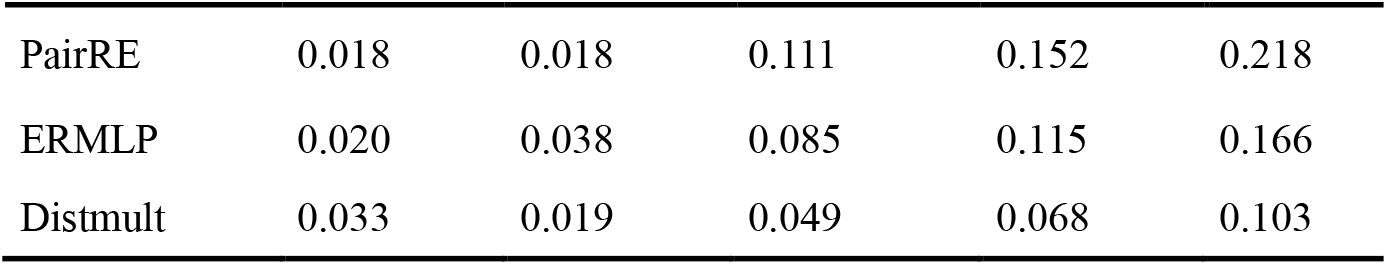
Internal Evaluation Results of 3 Models.

#### 3.4.2 External evaluation result

PairRE demonstrated the fastest first hit and the best speed of hitting half of the KP clinical trial drugs overall in the external validation as seen in Table 6, suggesting that it excels at making predictions quickly and consistently. In contrast, ERMLP was slower on first hits and hitting half of the KP clinical trial drugs, but ended up hitting all of the clinical trial drugs in an earlier position, which may reflect the fact that the ERMLP generalization Limited ability to deal with the full range of correct predictions more centrally, covering most of the correct predictions more quickly rather than extending to more samples, may mean that it performs poorly when it encounters more difficult predictions.

**Table 6.** External Evaluation Results of 3 models.

|  | First_hit | Median_hit | Last_hit |
| --- | --- | --- | --- |
| PairRE | 102 | 944 | 7947 |
| ERMLP | 305 | 1592 | 5893 |
| Distmult | 238 | 2400 | 13188 |

## 4 Conclusion

### 4.1 Conclusion

This investigation identified five potential drug candidates to combat the KP virus using deep learning methods. Key findings include:

1. **Drug Selection**: From the DrugBank database, 3,475 drugs of biotech and small-molecule types were selected, excluding compounds containing seleninic acids, bad-odor propionic acids, and radio-trace agents.
2. **Model Accuracy**: The CNN-DNN model achieved a validation accuracy of approximately 72%.
3. **Similarity Analysis**: Using ESM and cosine similarity analysis, five promising drug candidates with 85% similarity to KP virus strains were identified.
4. **Predictive Model Performance**: The DRKG approach demonstrated that PairRE consistently ranks correct drugs higher and outperforms other models across multiple evaluation metrics, making it the most effective predictive method.

In summary, this study presents a rapid and efficient framework for drug discovery, leveraging deep learning (CNN-DDIs) for validation and ESM with cosine similarity analysis to identify promising drug candidates. The proposed method reduces drug discovery cycle time to mere months, compared to the extensive timelines required by traditional laboratory-based approaches, offering a scalable and effective solution for targeting bacterial pathogens like KP.

### 4.2 Remarks

This paper presents a method for data extraction, followed by CNN/DDIs and an ESM model with cosine similarity analysis, to identify promising drugs for combating KP. These drug candidates will undergo rigorous preclinical and clinical trials to validate their efficacy against KP infections. Additionally, an ongoing study compares known drugs used to treat KP with the selected 3,475 biotech and small-molecule drugs through cosine similarity analysis, aiming to cross-examine and identify potential candidates for repurposing, if they differ.

## Declarations

## Ethics approval and consent to participate

Not Applicable.

## Consent for publication

Not Applicable.

## Availability of data and materials

The datasets and computer codes supporting this study are publicly available at the following repository: https://github.com/Peng-Bio/DL-Drugs-KP/upload/main.

## Authors’ contributions

P. L., J. Z., S. M.: Performed data analyses, constructed the model, and drafted the manuscript. Z. L.: Responsible for manuscript drafting and refinement. L. J.: Curated data and performed sequence analyses. H. Q.: Conducted the project and drafted the manuscript. Y. X., T. H.: Constructed the model and generated downstream analyses. Y. B.: Provided clinical advice and drafted manuscript. G. Q., W. Z.: Conducted the project, conceived the study, and revised the manuscript.

## Acknowledgements

This research was supported by a Grant from the Innovation Fund of Jinying Technology Co., Ltd. (grant no. CX: 20210506D) and a Grant from the Hong Kong Innovation and Technology Commission (grant no. PRP/062/22FX). The authors would like to express their heartfelt gratitude to Jinying Technology Co., Ltd. for their generous funding and continuous support, which made this study possible. We also thank all collaborators and research teams for their invaluable contributions to this work.

